# Bi-directional encoding of context-based odor signals and behavioral states by the nucleus of the lateral olfactory tract neurons

**DOI:** 10.1101/2020.07.30.230060

**Authors:** Yuta Tanisumi, Kazuki Shiotani, Junya Hirokawa, Yoshio Sakurai, Hiroyuki Manabe

## Abstract

The nucleus of the lateral olfactory tract (nLOT) is not only a part of the olfactory cortex that receives olfactory sensory inputs from the olfactory bulb, but also one of the cortical amygdala areas that regulates motivational behaviors. To examine how the neural ensemble activity of the nLOT is modulated by motivational processes that occur during various states of learned goal-directed behaviors, we recorded nLOT spike activities of mice performing odor-guided go/no-go tasks for obtaining a water reward. We found that the majority of the nLOT neurons exhibited sharp go-cue excitation and persistent no-go-cue inhibition responses triggered by an odor onset. The bi-directional cue encoding introduced nLOT population response dynamics and provided a high odor decoding accuracy before executing cue-odor-evoked behaviors. The go-cue preferred neurons were also activated in the reward drinking state, indicating context-based odor-outcome associations. These findings suggest that the nLOT neurons play an important role in the translation from context-based odor information to appropriate behavioral motivation.

## Introduction

The nucleus of the lateral olfactory tract (nLOT) is a part of the olfactory cortex that receives direct sensory inputs from the olfactory bulb and the olfactory cortex, such as the piriform cortex (*1, 2*). Alternately, it also receives projections from the anterior amygdaloid area, anterior cortical and posterolateral cortical amygdaloid nuclei, and amygdalo-piriform transition area, and forms part of the olfactory amygdala (*3*). While some authors have considered the nLOT as a component of the olfactory cortex (*1, 4*), others have regarded it as a component of the cortical amygdala areas that plays a critical role in generating odor-driven behaviors (*5*). The nLOT not only has bi-directional connection with the olfactory bulb and piriform cortex, but also strongly innervates the basolateral amygdala and ventral striatum (*1, 2, 6*). Due to its anatomical features, it is possible that the nLOT is involved in odor-evoked motivational behaviors.

In addition to these anatomical evidences, a recent study (*7*) showed functional evidence that nLOT integrity was required for the normal functioning of the olfactory system. The researchers conducted a series of behavioral tests using rats that were submitted to bilateral excitotoxic lesions of the nLOT. The nLOT-lesioned rats exhibited severe olfactory deficits with an inability to detect and discriminate between odors.

Despite the accumulation of knowledge, there are no reports of the *in vivo* recording of neuronal activity in the nLOT; therefore, the electrophysiological features of the nLOT neurons on odor-evoked motivational behavior have not been clarified. The purpose of our study was to investigate how the neural activity was modulated by motivational processes that occurred during various behavioral states in a goal-directed task. Here, we recorded the neural spike activities in the nLOT of freely moving mice performing an odor-guided go/no-go task. We found that the majority of nLOT neurons exhibited go-cue excitation and no-go-cue inhibition responses triggered by an odor onset. The bi-directional cue encoding strongly contributed to the nLOT neuron population dynamics before executing cue-odor-evoked behaviors; additionally, the go-cue preferred neurons encoded reward drinking state, indicating context-based odor-outcome associations. Our results suggest that the nLOT is critical for encoding context-based cue-outcome signals, and may play an important role in the translation of odor stimulus information to odor-guided behavioral motivation.

## Results

We recorded from 365 well-isolated neurons in the nLOT of four mice performing odor-guided go/no-go tasks (Figs. 1A and 1B). Briefly, the go trial required the mice to first sample a go-cue odor stimulus presented at an odor port and then to move to a reward port for obtaining a water reward. Conversely, the no-go trial required the mice to first sample a no-go-cue odor stimulus presented at the odor port and then to stay near it to wait for the next trial. It is important to note that the mice were required to keep their nose inserted into the odor port during odor presentation (500 msec). For all mice, the median of the odor-sampling epoch (the time from the odor valve opening until the withdrawal of the snout by the mouse from the odor port) was 788 msec (interquartile range: 669–962 msec) in the go trials, and 642 msec (interquartile range: 562–798 msec) in the no-go trials (44 sessions from four mice). In the following sections, we describe our analyses of the neural activity recorded during odor-sampling and the following odor-guided behaviors.

**Fig. 1.**
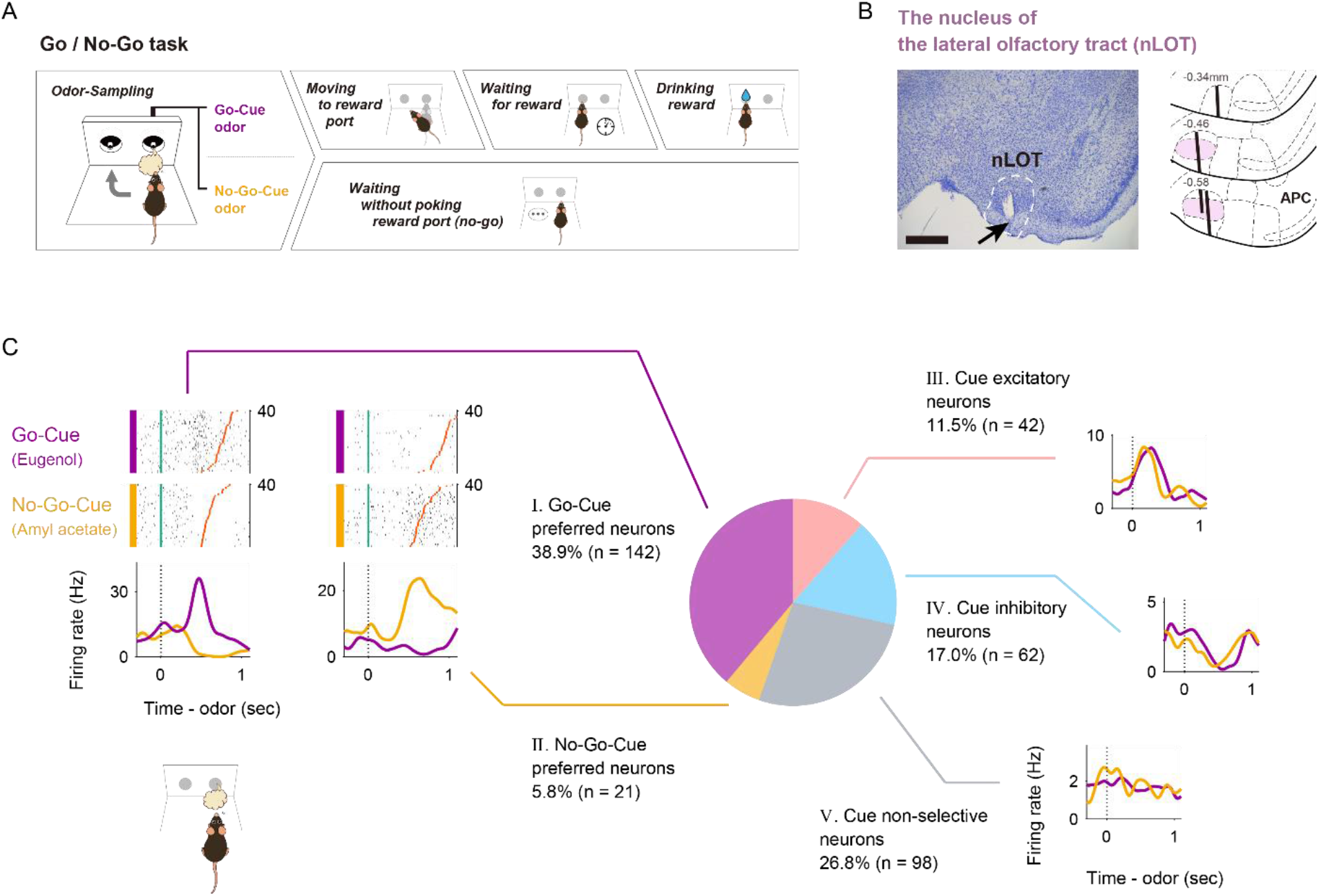
Nucleus of the lateral olfactory tract (nLOT) neuron activity patterns during the odor-guided go/no-go task. (A) Time course of the odor-guided go/no-go task. Behavioral epoch temporal progression from left to right. (B) Nissl-stained frontal section (an arrow indicates a tip of the tetrodes) and recording tracks (vertical thick lines) of the nLOT. The pink areas show layer II of the nLOT. APC, anterior piriform cortex. Scale bar: 500 μm. (C) Example firing patterns of nLOT neurons during the odor-sampling epoch (the time from the odor valve opening to odor port exit) in the odor-guided go/no-go task. Each row contains the spikes (black ticks) for one trial, aligned to the time of odor valve opening (corresponding to the odor port entry, green ticks). Red ticks refer to the times of odor port exit. The correct trials are grouped by odor and within each group are sorted by the duration of the odor-sampling epoch (40 selected trials from the end of the session are shown per category). Histograms are averaged across odors and calculated using a 20 msec bin width and smoothed by convolving spike trains with a 60 msec-wide Gaussian filter (purple, go-cue odor; orange, no-go-cue odor). Vertical dashed lines indicate the time of the odor valve opening. nLOT neurons were classified into five types (purple pie, type I; orange pie, type II; pink pie, type III; light blue pie, type IV; and gray pie, type V) based on the odor-sampling epoch response (Fig. S1).

### The five-type classification of nLOT neurons based on the odor-sampling epoch response

As the nLOT receives direct inputs from the mitral cells of the olfactory bulb, we first focused on the neural activity during the odor-sampling epoch. We found that the firing rates of the nLOT neurons increased or decreased during the odor-sampling epoch. For a large subset of the neurons, these firing rate changes depended on whether the presented odor was go-cue or no-go-cue (examples shown in Fig. 1C, left). In order to quantify the dependence of the firing rate on cue odor presentation, we used a receiver operating characteristic (ROC) analysis approach. We calculated the firing rate changes from baseline (1,000 to 0 msec before the end of the inter-trial interval) during the odor-sampling epoch. Across the population, 73.2% of the nLOT neurons exhibited significant responses for at least one of the cue odor presentations (Fig. S1, p < 0.01, permutation test). In this cue odor selective population, we also calculated the preference for go-cue and no-go-cue odor presentation. We observed that 53.2% of the population showed a significant go-cue odor preference, whereas 7.9% of them showed a significant no-go-cue odor preference (Fig. S1, p < 0.01, permutation test). The other population exhibited increased or decreased responses to both go-cue and no-go-cue odor presentations. Thus, most of the nLOT neurons showed a wide variety of firing rate changes during an odor-sampling epoch.

Based on these response profiles of odor-sampling epoch, we classified the nLOT neurons into five types (Figs. 1C and S1, Table 1, see Materials and methods). The first neuron group (type I, 38.9% of all neurons) exhibited significant preference for the presented go-cue odor––we will refer to these as “go-cue preferred neurons” (purple pie chart in Figs. 1C and S1). The second neuron group (type II, 5.8% of all neurons) exhibited significant preference for the presented no-go-cue odor––we will refer to these as “no-go-cue preferred neurons” (orange pie chart in Figs. 1C and S1). Two other neuron groups (type III and IV, 11.5% and 17.0% of all neurons) showed significant excitatory and inhibitory responses, respectively, for both presented cue odors without preference for a particular cue odor; we will refer to these as “cue excitatory neurons” (pink pie chart in Figs. 1C and S1) and “cue inhibitory neurons” (light blue pie chart in Figs. 1C and S1), respectively. The remaining neuron group (type V, 26.8% of all neurons) did not show significant responses for either presented cue odors––we will refer to these as “cue non-selective neurons” (gray pie chart in Figs. 1C and S1). This classification demonstrated the diverse cue encoding patterns in the nLOT, suggesting that the nLOT neurons do not represent a particular odorant profile from the olfactory bulb, and instead represent the complex and diverse odor information leading to odor-guided behaviors.

**Table 1.** The distribution of types of nucleus of the lateral olfactory tract neurons

| Mouse | I. Go-Cue<br>preferred<br>neurons | II. No-Go-Cue<br>preferred<br>neurons | III. Cue<br>excitatory<br>neurons | IV. Cue<br>inhibitory<br>neurons | V. Cue<br>non-selective<br>neurons | (Total) |
| --- | --- | --- | --- | --- | --- | --- |
| #1 | 8 | 5 | 10 | 21 | 28 | 72 |
| #2 | 8 | 4 | 3 | 9 | 16 | 40 |
| #3 | 22 | 5 | 12 | 31 | 31 | 101 |
| #4 | 104 | 7 | 17 | 1 | 23 | 152 |
| (Total) | 142 | 21 | 42 | 62 | 98 | 365 |

### Go-Cue preferred neurons bi-directionally encode cue odors with excitations and inhibitions

Among the go-cue preferred neurons (type I neurons, n = 142), which represent the major population of the nLOT neurons (Fig. 1C), each neuron showed a sharp peak in the firing rate after ~600 msec of go-cue odor presentation and persistent inhibition during the latter part of the no-go-cue odor sampling epoch (Fig. 2A). To quantify the dynamics of this bi-directional cue encoding, we calculated the firing rate changes from the baseline (200 to 0 msec before the end of the inter trial interval) in the sliding bins during the odor-sampling epoch for each neuron. For each accurate trial type, we calculated the area under the ROC curve (auROC) value at each time bin (width: 100 msec, step: 20 msec) (Figs. 2B and 2C), and three measures from the auROC values: “onset time,” “time of center of mass,” and “duration” (Fig. 2D, see Materials and methods). The onset times of the go-cue excitations were earlier than those of no-go-cue inhibitions (p < 10^-4^, Wilcoxon rank-sum test). Regarding the times of the center of mass and durations, the go-cue excitation responses were earlier (p < 10^-23^, Wilcoxon rank-sum test) and sharper (p < 0.05, Wilcoxon rank-sum test) than the no-go-cue inhibition responses. Conversely, the no-go-cue inhibition responses were sustained until the mice withdrew their snouts from the odor port. For each neuron, both the go-cue excitation response and the no-go-cue inhibition response were significantly observed particularly 450–550 msec after the odor onset (Figs 2E and S2, p < 0.01, permutation test). Thus, each go-cue preferred neuron exhibited both a sharp go-cue excitation and a persistent no-go-cue inhibition at the specific times during the odor-sampling epoch.

**Fig. 2.**
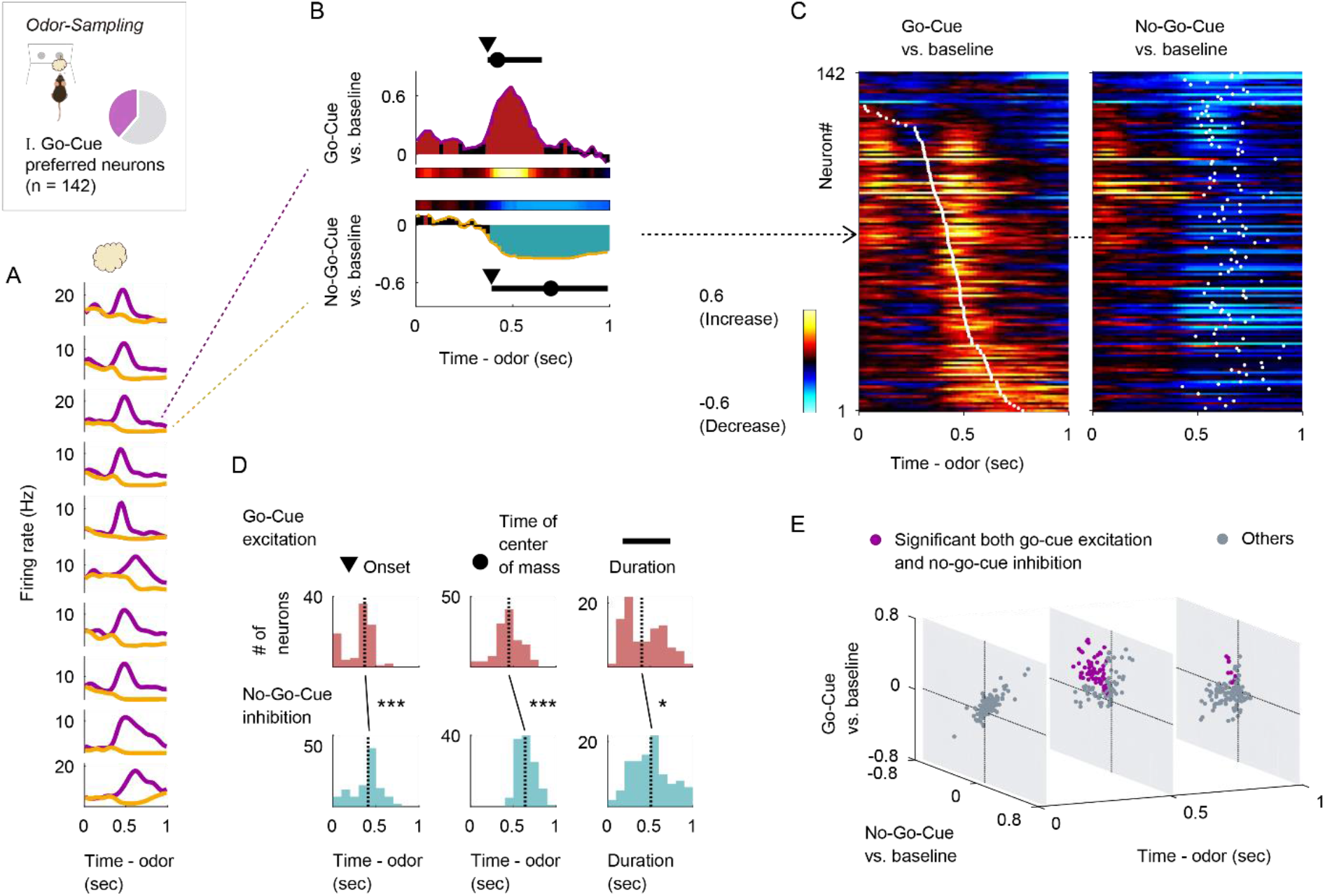
Go-Cue preferred neurons showed phasic excitation to go-cue odor and persistent inhibition to no-go-cue odor. (A) The example firing patterns of go-cue preferred neurons during the odor-sampling epoch. Spike histograms are calculated using a 20 msec bin width and smoothed by convolving spike trains with a 60 msec wide Gaussian filter (purple line, go-cue odor; orange line, no-go-cue odor). (B) Example of the area under the receiver operating characteristic curve (auROC) values for a go-cue preferred neuron. The auROC values (aligned by odor valve opening) were calculated by go-cue odor presentation versus baseline (top) and no-go-cue odor presentation versus baseline (bottom) in the sliding bins (width, 100 msec; step, 20 msec). The red bars show significant excitation and blue bars show significant inhibition (p < 0.01, permutation test). Based on the significant time points, onset time (black triangle), time of center of mass (black circle), and duration (black horizontal line) were calculated. (C) The auROC values for go-cue preferred neurons (n = 142, type I neurons). Each row corresponds to one neuron, with neurons in the left and right graphs in the same order. The neurons are sorted by the times of center of mass (white dots) of the auROC values calculated by go-cue odor presentation versus baseline. The color scale is as in (B). An arrow indicates the same neuron as in (B). (D) Distributions of onset time, time of center of mass, and duration for significant excitations (top, red) and significant inhibitions (bottom, blue). Vertical dashed lines indicate median values. Statistical significance between excitations and inhibitions (*p < 0.05, ***p < 0.001) was assessed by the Wilcoxon rank-sum test. (E) Time course of excitation to go-cue odor and inhibition to no-go-cue odor. Purple dots, significant both go-cue excitation and no-go-cue inhibition (p < 0.01, permutation test); gray dots, other responses.

It is possible that the sharp go-cue excitation responses correlated with the executions of the go behaviors. To verify this possibility, we compared the peak firing rates and the half width of firing in the go-cue excitation between the two alignment conditions (odor valve opening versus odor port exit). We observed that the peak firing rates were higher relative to the odor onset (p < 10^-15^, Wilcoxon signed-rank test) and the temporal organizations were significantly tighter (p < 0.001, Wilcoxon signed-rank test) than the firing rate relative to the odor port exit (Figs. S3A-B). These results indicated that the go-cue excitation responses of the go-cue preferred neurons were triggered by odor onset rather than execution of the go behavior. Furthermore, the distinct cue responses were observed in the correct go trials, and not in the trials that were correct no-go, error, or odorless (Fig. S3C), suggesting that the distinct go-cue excitation responses reflected signals eliciting appropriate motivational behavior. Notably, the intensities of the majority of the cue responses were kept stable across trials (Fig. S3D). In conclusion, the distinct go-cue excitation responses were triggered by odor onset and stable with respect to the appropriate odor-guided behaviors.

### nLOT neuron population exhibits rapid response dynamics before executing cue-odor-evoked behaviors

We demonstrated that the go-cue preferred neurons exhibited specific temporal dynamics during odor-sampling as a representative population of nLOT neurons (Fig. 2). Similarly, the no-go-cue preferred neurons exhibited both no-go-cue excitation and persistent go-cue inhibition during cue odor presentation at ~600 msec (Fig. S4). The cue excitatory neurons and the cue inhibitory neurons also changed their firing rates during odor-sampling (Figs. S5 and S6). Thus, the nLOT neurons exhibited diverse firing patterns and complex temporal dynamics during odor-sampling. In this section, we examined the nLOT population encoding and the contribution of each neuron group during odor-sampling using different methods of analysis.

Calculating go-cue versus no-go-cue preference during odor-sampling clearly showed the strong encodings of cue preference at 400–500 msec after the odor onset across the population (Fig. 3A, p < 0.01, permutation test). To gain insight into the dynamics of the population response, we visualized the average population activity using principal component analysis, a dimensionality reduction method (Fig. S7). Fig. 3B shows trajectories of the mean response of the nLOT neuron population to go-cue and no-go-cue odors, represented as the projections onto the first three principal components (PC) during the odor-sampling epoch. Throughout the approximately 300 msec interval from the odor onset, trajectories remained converged, showing little difference across conditions. Over the late phase of odor-sampling, specifically 400–500 msec from the odor onset, trajectories in the odor-sampling epoch subspace began to spread out and clearly separated at the population level. To quantify these observations, we measured the instantaneous separation between the population cue responses (Fig. 3C). The separation reached a maximum at ~500 msec and remained above the baseline levels until the odor port exit. Additionally, we calculated the rate at which the population activity vectors changed (width: 100 msec, step: 20 msec; Fig. 3D). These rates increased to a maximum within ~500 msec and remained above the baseline levels over the initiations of cue-odor-evoked behaviors (go or no-go behaviors). Thus, the nLOT neuron population dramatically showed profound transformations in the dynamics of cue encoding at 400– 500 msec after the odor onset.

**Fig. 3.**
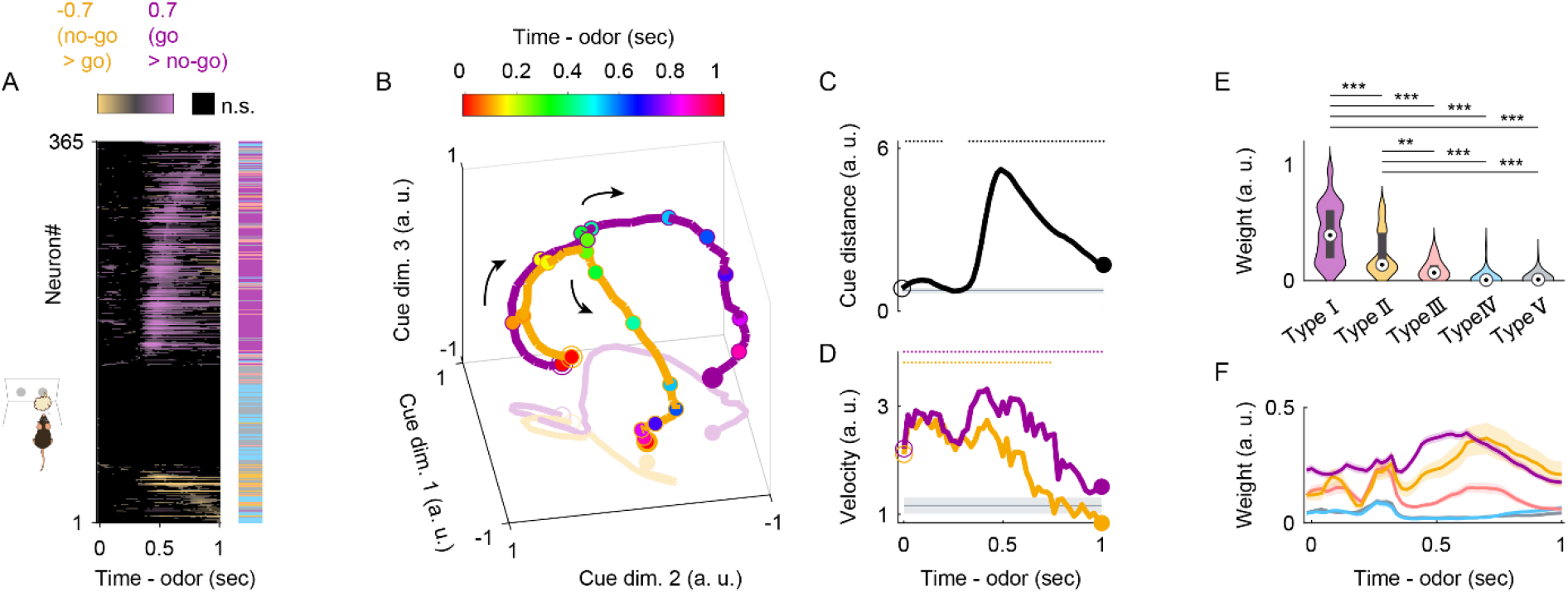
nLOT neuron population response before the initiation of odor-guided behaviors. (A) The auROC values (go-cue versus no-go-cue odor presentation, aligned by odor valve opening) for all neurons. Each row corresponds to one neuron. Neurons are sorted by the peak time for the auROC values. The color scale indicates significant preferences (p < 0.01, permutation test; positive values correspond to the go-cue preferred responses). The black boxes indicate bins with non-significant preferences (p > 0.01, permutation test). The colored box on the right shows the neuron type for each neuron (purple, type I; orange, type II; pink, type III; light blue, type IV; gray, type V). (B) Visualization of the nLOT neuron population responses during odor-sampling epoch using principal component analysis (n = 365 nLOT neurons). The responses to cue odors are projected onto the first three principal components corresponding to the odor-sampling epoch subspaces. Purple line, go-cue odor; orange line, no-go-cue odor. Temporal progression is depicted from unfilled purple/orange spheres to filled purple/orange spheres. (C) The distance between nLOT neuron population responses. The gray line and shaded areas show the mean ± 2 standard deviation (SD) baseline values during the baseline epoch. Top dots indicate the time bins showing values more than mean + 2 SD baseline values. (D) Rate of change (velocity) of nLOT neuron population responses. Purple line, go-cue odor; orange line, no-go-cue odor. The gray line and shaded areas show the mean ± 2 SD baseline values during the baseline epoch. Top dots indicate the time bins showing values more than mean + 2 SD baseline values. (E) Neural weights in the first dimension of the odor-sampling epoch subspaces. Box-Plots in violin-plots indicate medians and interquartile ranges. Purple, type I; orange, type II; pink, type III; light blue, type IV; gray, type V. The statistical significance among five groups (**p < 0.01, ***p < 0.001) was assessed by a one-way analysis of variance (ANOVA) with Tukey’s post hoc test. (F) Neural weights along the time course in the first dimension of each sliding bin (width: 100 msec, step: 20 msec). The shaded areas represent ± SEM. Purple, type I; orange, type II; pink, type III; light blue, type IV; gray, type V.

Next, we examined the mechanism of the contribution of individual nLOT neurons to the population response to evaluate the absolute values of PC coefficients as the neural weights (Figs. 3E and S7A-C). The values of the neural weights in the first dimension of the odor-sampling epoch subspaces showed that type I neurons contributed considerably to the population response. To further examine the contributions along the time course, we calculated the absolute values of the PC coefficients in the sliding bins (width: 100 msec, step: 20 msec) during the odor-sampling epoch (Figs. 3F and S7D). The values of the neural weights in the first dimension of each bin exhibited significant contributions of type I neurons to the population response, especially during 400–500 msec after the odor onset, corresponding to the dynamics of cue encoding. These results indicated that the go-cue preferred neurons strongly contributed to the profound transformations in the dynamics of nLOT cue encoding.

### nLOT neurons provided sufficient information to account for behavioral accuracy

To examine whether the population activity accounted for the animals’ behavioral accuracy, we performed a decoding analysis. This analysis determined whether the firing rates of the nLOT neuron populations could be used to classify each individual trial as go or no-go. We used support vector machines with linear kernels as a decoder. Analyses of the decoding time course based on nLOT neurons using a sliding time window revealed that the decoding accuracy was maintained at chance levels 300 msec after the odor onset and subsequently dramatically increased above the level of animals’ behavioral accuracy 400–500 msec after the odor onset (Fig. 4A). In the 400–500 msec period, 124 neurons provided sufficient information to account for the behavioral accuracy (the top right panel in Fig. 4A). Thus, fewer than 150 nLOT neurons provided sufficient information to account for behavioral accuracy at least 500 msec after the odor onset.

**Fig. 4.**
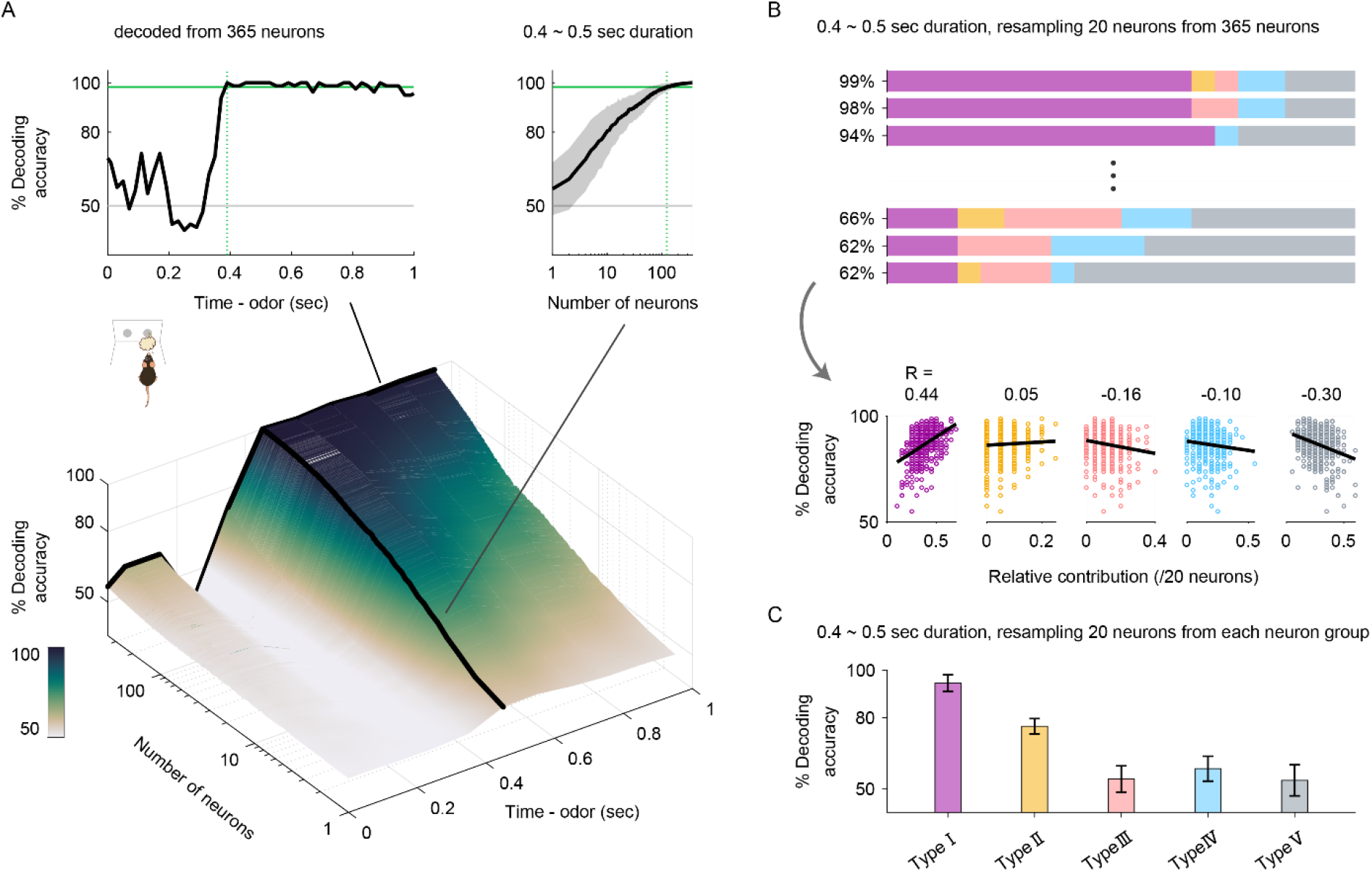
nLOT neurons provided sufficient information to account for behavioral choices. (A) The time course of odor decoding accuracy. A vector consisting of instantaneous spike counts for 1–365 neurons in a sliding window (width, 100 msec; step, 20 msec) was used as an input for the classifier. Training of the classifier and testing were performed at every time point. Green horizontal lines indicate the level of animal behavioral performance. Gray horizontal lines indicate the chance level (50%). Green vertical dashed lines indicate the first points wherein the decoding accuracy reached the level of the animal behavioral performance. The areas with shading represent ± SD. (B) Decoding accuracies based on 20 randomly sampled neurons without replacement during the 400–500 msec period after the odor onset. Bottom, correlation between the decoding accuracy and relative contribution as the proportion of each neuron type number in randomly sampled neurons. Purple, type I; orange, type II; pink, type III; light blue, type IV; gray, type V. (C) Decoding accuracies based on 20 randomly sampled neurons from each nLOT neuron group without replacement during the 400-500 msec period after the odor onset. The error bars represent ± SD. Purple, type I; orange, type II; pink, type III; light blue, type IV; gray, type V.

To further examine the contribution of neural decoding for each nLOT neuron group, we calculated the decoding accuracy based on 20 randomly sampled neurons without replacement during the 400–500 msec period after the odor onset (Fig. 4B). The proportions of type I neurons in the sampled datasets were correlated with the decoding accuracies (Figs. 4B and S8). Next, we sought to rule out the possibility that the bias of the neuron group size affected the contribution of neural decoding. To rule out this prospect, we also calculated the decoding accuracy based on 20 randomly sampled neurons from each nLOT neuron group (Fig. 4C). The decoding accuracy using the type I neuron group reached more than 90%, indicating that only a small number of type I neurons provided more information to account for behavioral choices. Thus, the nLOT had a large population of neurons that strongly correlated with the animals’ behavioral choices.

### Bi-Directional cue-outcome encoding following odor-guided behaviors

Our analyses of the dynamics of cue encoding suggest that many nLOT neurons maintained cue selective responses during cue-odor-evoked behaviors after odor-sampling. Notably, the persistent inhibition responses of type I neurons during the no-go-cue odor-sampling were sustained over the odor port exit (Figs. 2B-D). This raises the query of whether the selectivity disappears or persists once the cue-odor-evoked behaviors are executed. By aligning the neural activity to behavioral events (event-aligned spike histograms, see Materials and methods), we noticed that type I neurons were selective for the reward drinking behavior after go-cue odor-sampling and the persistent inhibition responses were sustained during the no-go waiting behavior after no-go-cue odor-sampling (Fig. 5A). We quantified the response profiles of each neuron group during odor-evoked behaviors by calculating the firing rate changes from baseline (Figs. 5B-C and S9A, the spike data were aligned to the odor port exit and water port entry). Across the population, many type I neurons showed significant excitatory responses for the drinking behavior (purple histogram at the top in Fig. 5B, p < 0.01, permutation test), and significant inhibitory responses for the no-go waiting behavior (purple histogram at the bottom in Fig. 5C, p < 0.01, permutation test). The drinking responses of type I neurons were higher than those of other groups and the no-go waiting responses of type I neurons were lower than those of other groups, indicating that they were type I neuron specific responses (Fig. S9B, one-way analysis of variance with Tukey’s post hoc test). For each neuron, the inhibitions were maintained for 800 msec (interquartile range: 290–1480 msec) from the initiation of the no-go behavior. Thus, type I neurons exhibited associations between the go-cue excitations and excitatory responses for drinking behavior with persistent no-go-cue inhibitions, suggesting that nLOT neurons are involved in cue-outcome associations.

**Fig. 5.**
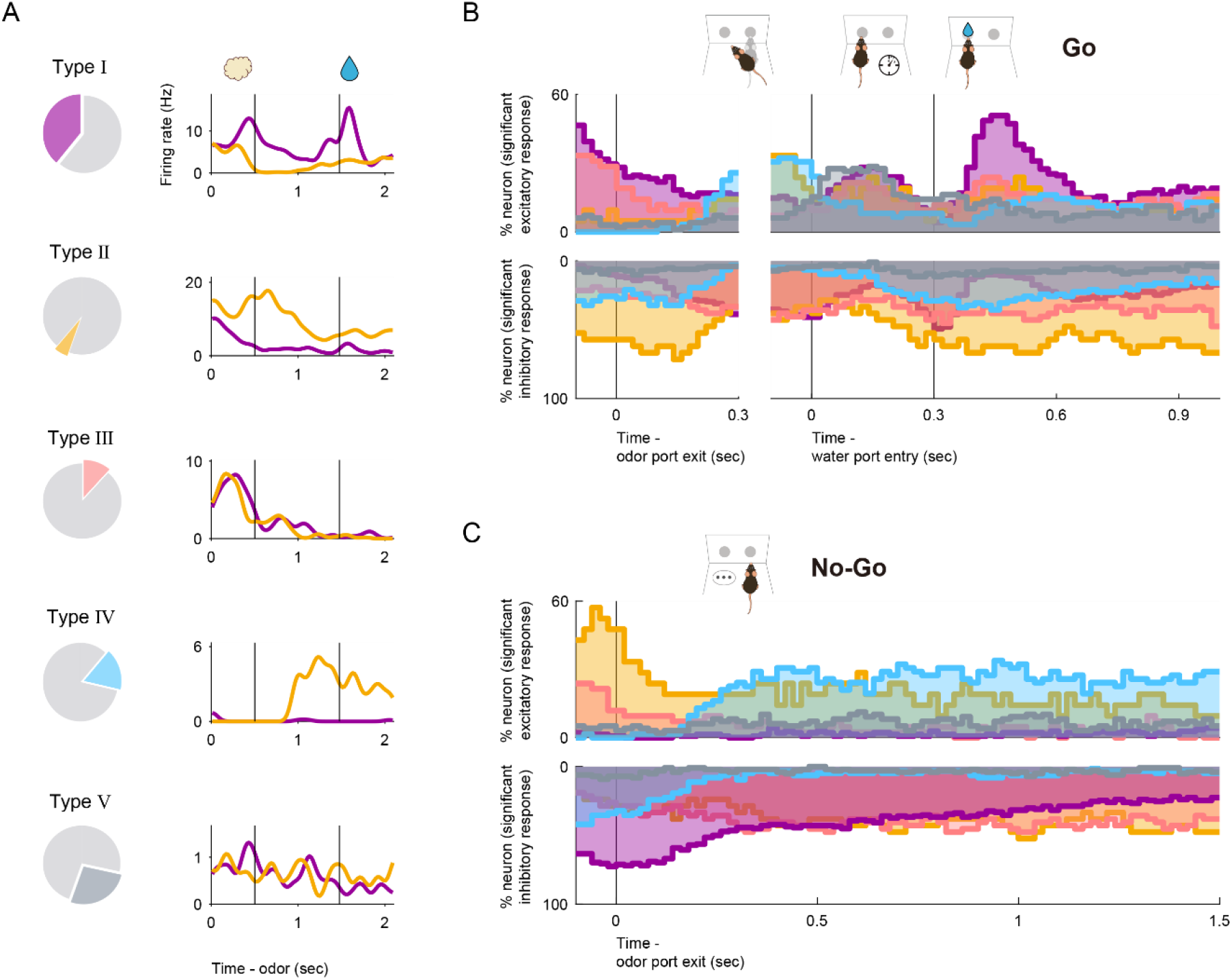
nLOT neurons exhibits bi-directional cue-outcome encoding following odor-guided behaviors. (A) An example firing pattern of each neuron group following odor-guided behaviors. Event-aligned spike histograms are calculated using a 20 msec bin width and smoothed by convolving spike trains with a 60 msec wide Gaussian filter (purple line, go trial; orange line, no-go trial). Vertical black lines indicate the odor valve offset and the onset of the water reward. (B) The proportions of neurons that exhibited significant responses were calculated from the auROC values (p < 0.01, permutation test) in go correct trials for each neuron group (top, excitation; bottom, inhibition). Vertical black lines indicate the odor port exit, water port entry, and the onset of the water reward. Purple, type I; orange, type II; pink, type III; light blue, type IV; gray, type V. (C) Same as (B), for no-go correct trials.

We aimed to determine if the other neuron groups responded to the cue-odor-evoked behavioral states. Many type II neurons showed significant inhibitory responses for the drinking behavior (orange histogram at the bottom in Fig. 5B, p < 0.01, permutation test) and significant excitatory responses for the no-go waiting behavior (orange histogram at the top in Fig. 5C, p < 0.01, permutation test). The drinking responses of type II neurons were lower than those of type I neurons and the no-go waiting responses of type II neurons were higher than those of type I neurons (Fig. S9B, one-way analysis of variance with Tukey’s post hoc test). Thus, the type I and type II neurons contrastingly encoded the go/no-go behavioral states after the odor-sampling. Furthermore, a subset of type III, IV, and V neurons tended to show an excitatory response in a specific time window in behavioral epochs, with inhibitory responses relative to other behavioral epochs (Fig. S9C). Particularly, each type IV neuron maintained the excitatory response to the no-go waiting state (light blue histogram at the top in Fig. 5C, p < 0.01, permutation test) for 560 msec (interquartile range: 205–1200 msec) from 290 msec (interquartile range: 210–650 msec) after initiation of the no-go behavior. These results indicate that each nLOT neuron group showed a specific firing pattern during odor-guided behaviors, depending on the response profiles in the odor-sampling epochs.

## Discussion

### Electrophysiological features of the nLOT neurons

The purpose of the study was to understand the electrophysiological features of nLOT neurons on motivational processes that occur during various behavioral states in odor-guided go/no-go tasks. In this study, we provided the first recording of neuronal activity in the nLOT in freely behaving mice performing odor-guided go/no-go tasks. Indeed, nLOT neurons exhibited diverse neural activities in response to odor presentations and cue odor-evoked behaviors in the task.

Previous anatomical studies have shown that the nLOT receives odor information from the olfactory bulb and various areas of the olfactory cortex, including the piriform cortex (*1, 2*). Subsequently, nLOT neurons project to the ventral striatum consisting of the olfactory tubercle (OT) and the nucleus accumbens (NAc) and also sends the axon into the basolateral amygdala (BLA) (*1, 2, 6*) that plays the critical role of regulating motivated behaviors (*8–10*). Moreover, a recent study (*7*) showed that nLOT-lesioned rats exhibited olfactory-related behavioral deficits with an incapacity to identify and discriminate between odors and interfered with the display of innate odor-evoked behaviors, such as sexual behavior, aggression, and avoidance of predators. Despite the accumulation of knowledge, the role of nLOT in the functional circuit to convert odor information into appropriate behaviors have not been clarified.

In this study, we classified five types of neurons on the basis of their firing pattern during the odor-sampling epochs. A majority of nLOT neurons (type I neurons, go-cue preferred neurons) exhibited phasic excitatory responses during go-cue odor-sampling epochs and sustained inhibitory responses during no-go-cue odor-sampling epochs (Figs. 1C and 2). The activity pattern of the no-go-cue preferred neurons (type II neurons) was opposite to that of the go-cue preferred neurons (Fig. S4). These bi-directional cue encoding patterns are similar to the cue encoding in the brain reward circuit, including the ventral striatum and the ventral tegmental area (*11–13*) rather than the olfactory circuit (*14–16*) during cue-outcome association tasks. We also demonstrated that the go-cue and no-go-cue preferred neurons highly contributed to the population dynamics of cue encoding and the decoding, for accuracy of the animal’s choices (Figs. 3 and 4), suggesting that these bi-directional response neurons for cue odors effectively provide sufficient information to account for behavioral choices. These bi-directional cue encoding small number of neurons having high level of information may be effective in the nLOT with only a small volume of 0.24 mm^3^ and 19,000 neurons (*17*).

The go-cue preferred neurons also showed firing activities during drinking behavior (Fig. 5B), consistent with other brain areas involved in motivational processes (*18, 19*). Additionally, the go-cue preferred neurons exhibited suppressed responses to the no-go waiting behavioral states (Figs. 5C and S9B). Moreover, the no-go-cue preferred neurons suppressed their firing activity during the go behavioral states and exhibited excitatory activities in the no-go behavioral states (Figs. 5B-C and S9B). These results suggest that these nLOT neurons functionally associate the cue odor with the precise task outcomes derived from the odor information. Additionally, since the decoding analysis revealed that the responses of the go-cue and no-go-cue preferred neurons during the odor-sampling epoch contained information dictating the animal’s choice (Fig. 4), we speculate that the nLOT is one of the critical components of the circuitry responsible for creating and providing signals eliciting appropriate motivational behavior into the motivation circuits, including the ventral striatum and BLA. Due to the function of nLOT, we assume that the lesion of nLOT caused inhibition of olfactory-driven behaviors (*7*).

The subsets of type III, IV, and V neurons exhibited an excitatory response in a specific time window in behavioral epochs with inhibitory responses relative to other behavioral epochs (Fig. S9C). These results raise the probability that they represent the behavioral context of the task. A recent study showed the brain-wide global representation of the state-dependent activity during odor-guided motivated behavior (*20*). We assume that the context-dependent activities of the type III, IV, and V neurons may contribute to the brain-wide specific information processing mode in the brain.

### Neural circuits including the nLOT

Olfactory information is transmitted into the nLOT with a three-layered structure (layers I, II, and III). The nLOT layer II neurons contributing to over 80% of the total neuronal population of the nLOT (*17*), project to the dwarf cell regions in the OT (*6*). The OT sends a major projection to the ventral pallidum regulating expected positive and negative valences (*21–23*). The layer II neurons also project to the NAc shell (*6*) that processes hedonic or motivational value (*24–27*). Similar to the neural responses in these areas, we demonstrated the reward-predicting cue and reward signals of type I neurons in the nLOT (Figs. 2 and 5). In the context of a recent frontal cortex research showing a connectivity-defined neuron type that carries a single variable (*28*), we speculate that type I neuron outputs in the nLOT layer II to the OT and NAc contribute to the encoding of the positive or negative valences of expected and actual outcomes, and hedonic or motivational value. Conversely, the nLOT layer III neurons project to the BLA (*6*) that is an essential component of the amygdala underlying fear conditioning memory (*8, 29*). We demonstrated the no-go-cue preferred responses of type II neurons (Figs. 5 and S4) and the sustained positive responses to the no-go behaviors of type IV neurons. We assumed that these specific firing patterns in the nLOT layer III may contribute to the fear conditioning memory circuits. However, we have not verified the firing pattern of the nLOT neurons in the fear memory tasks. Future experiments are required to monitor the changes in the firing activity in the nLOT during odor-punishment association tasks.

We acknowledge that there are several limitations in this study. First, although we performed the first *in vivo* recording of neuronal activity in the nLOT during only an odor-guided go/no-go task, our data do not reflect the neuronal activity across different cue modalities, behavioral paradigms, and contexts. However, our data are potentially important in that the nLOT neural activity in freely behaving mice is modulated by the motivation of learned, odor-guided, and goal-directed behaviors, and may provide basic information regarding the nLOT encoding in positive and negative motivational contexts, reversal learning, and innate odor-driven behaviors (*30*). Second, a direct relationship between the distinct cue response of the nLOT neurons and context-dependent motivated behaviors is unclear. Third, the response profiles and functions of the nLOT specific projections to OT, NAc, and BLA on motivational processes have not yet been clarified. By using optogenetic manipulation or the fiber photometry tool to monitor cell-type and projection specific population activity, future studies can build on the paradigm and findings described here to address how the nLOT interacts with the projected areas to mediate the processes necessary for odor-guided behavior.

In conclusion, we extended the concept that nLOT integrity was required for the normal functioning of the olfactory system (*7*) and hypothesized that the nLOT plays a critical role in providing the odor information that elicited appropriate behavioral motivation into the motivation circuits in the odor-guided behavior. In a broad perspective, the verification of this hypothesis may have important implications for studying and leveraging neural circuits underlying odor-evoked motivation in health and disease.

## Materials and methods

### Animals

All the experiments were performed on male C57BL/6 mice (9 weeks old; weighing 20-25 g) purchased from Shimizu Laboratory Supplies Co., LTD., Kyoto, Japan. The mice were individually housed in a temperature-controlled environment with a 13-h light/11-h dark cycle (lights on at 8:00 and off at 21:00). They were provided with water after the training and recording sessions to ensure that the body weights dipped no lower than 85% of the initial levels and food was supplied ad libitum. All experiments were performed in accordance with the guidelines for animal experiments at Doshisha University and with the approval of the Doshisha University Animal Research Committee.

### Apparatus

We used a behavioral apparatus controlled by the Bpod State Machine r0.5 (Sanworks LLC, NY, USA), an open-source control device designed for behavioral tasks. The apparatus was comprised of a custom-designed mouse behavior box with two nose-poke ports on the front wall. The box was contained in another soundproof box (BrainScience Idea. Co., Ltd., Osaka, Japan) equipped with a ventilator fan that provided adequate air circulation and low-level background noise. Each of the two nose-poke ports had a white light-emitting diode (LED) and infrared photodiodes. Interruption of the infrared beam generated a transistor-transistor-logic (TTL) pulse, thus signaling the entry of the mouse head into the port. The odor delivery port was equipped with stainless steel tubing connected to a custom-made olfactometer (*31*). Eugenol was used as the go-cue odor and amyl acetate (Tokyo Chemical Industry Co., Ltd., Tokyo, Japan) as the no-go-cue odor. These odors were diluted to 10% in mineral oil and further diluted to 1:9 by airflow. Water reward delivery was based on gravitational flow, controlled by a solenoid valve (The Lee Company, CT, USA), and connected via Tygon tubing to stainless steel tubing. The reward amount (6 μL) was determined by the opening duration of the solenoid valve and was regularly calibrated.

### Odor-Guided go/no-go task

After a 3 sec inter-trial interval, each trial began by illuminating the LED light at the right odor port, which instructed the mouse to nose poke into that port. A nose poke into the odor port resulted in the delivery of one of the two cue odors for 500 msec. The mice were required to maintain their nose poke during odor stimulation to sniff the odor. After odor stimulation, the LED light was turned off and the mice could withdraw their noses from the odor ports. If eugenol odor (go-cue odor) was presented, the mice were required to move to and nose poke into the left water reward port within a timeout period of 2 sec. At the water port, the mice were required to maintain their nose poke for 300 msec before water delivery began. Next, 6 μL of water was delivered as a reward. If an amyl acetate odor (no-go-cue odor) was presented, the mice were required to avoid entering the water port for 2 sec following odor stimulation. Once in 10 trials, we introduced catch trials in which the air stream was delivered through a filter containing no odorants during which the mice were not rewarded whichever behavior they chose (go or no-go behavior). The accuracy rate was calculated as the total percentage of successes in the go and no-go trials in a session. The mice performed up to 448 trials (go error: ~20 trials, no-go error: ~4 trials, go in catch trials: ~11 trials, no-go in catch trials: ~37 trials) in each session per day.

### Electrophysiology

The mice were anesthetized with medetomidine (0.75 mg/kg i.p.), midazolam (4.0 mg/kg i.p.), and butorphanol (5.0 mg/kg i.p.), and implanted with a custom-built microdrive of four tetrodes in the nLOT (0.1 mm anterior to the bregma, 2.0 mm lateral to the midline). Individual tetrodes consisted of four twisted polyimide-coated tungsten wires (California Fine Wire, single wire diameter 12.5 μm, gold plated to less than 500 kΩ). Two additional screws were threaded into the bone above the cerebellum for reference. The electrodes were connected to an electrode interface board (EIB-18, Neuralynx, MT, USA) on the microdrive. The microdrive array was fixed to the skull with LOCTITE 454 (Henkel Corporation, Düsseldorf, Germany). After the completion of surgery, the mice received atipamezole (0.75 mg/kg i.p.) to reverse the effects of medetomidine and allow for a shorter recovery period. Additionally, the mice received analgesics (ketoprofen, 5 mg/kg, i.p.). Behavioral training resumed at least 1 week after the surgery. Electrical signals were obtained with an open-source hardware (Open Ephys). For unit recordings, signals were sampled at 30 kHz in Open Ephys and band-pass filtered at 600–6,000 Hz. After each recording, tetrodes were adjusted to obtain new units.

### Data analyses

All data analyses were performed using built-in and custom-built software in MATLAB 2019a (The Mathworks, Inc., MA, USA).

Spike sorting: Spikes were sorted into clusters offline based on their waveform energy, peak amplitudes, and the first principal components from the four tetrode channels using an automated spike-separation algorithm KlustaKwik (K.D. Harris). The resulting classifications were corrected and refined manually with the MClust software (A.D. Redish). The clusters were considered as single units only when the following criteria were met: (1) refractory period (2 msec) violations were less than 0.2% of all spikes and (2) the isolation distance, estimated as the distance from the center of the identified cluster to the nearest cluster based on the Mahalanobis distance, was more than 20.

Spike train analyses: Neural and behavioral data were synchronized by inputting each event timestamp from the Bpod behavioral control system into the electric signal recordings system. To calculate the firing rates during tasks, peri-event time histograms (PETHs) were calculated using a 20 msec bin width and smoothed by convolving spike trains with a 60 msec wide Gaussian filter.

To examine the relationship between the firing rate changes among individual nLOT neurons and the development of behavioral epochs in behavioral tasks, we created event-aligned spike histograms (EASHs) (*32*). As behavioral epoch durations varied for each trial, the median durations of the epoch were calculated first. In the odor-guided go/no-go task, the median duration of odor-sampling epochs (from the odor onset to the odor port exit) was 788 msec in the go trials, 642 msec in the no-go trials, and the median duration of moving epochs (from the odor port exit to the water port entry) was 388 msec. The spike timing during each epoch and for each trial was linearly transformed to correspond with the median behavioral duration of each epoch. The number of spikes in each epoch was preserved. Furthermore, we defined the waiting epoch (300 msec reward delay, from the water port entry to the onset of the water reward) and the drinking epoch (1,000 msec after the onset of the water reward). These epochs were not applied to the transformation because their durations did not change across trials. In this way, the regular raster plots were transformed into event-aligned raster plots. Consequently, an EASH was calculated using a 20 msec bin width and smoothed by convolving the spike trains with a 60 msec wide Gaussian filter from the event-aligned raster plots (Fig. 5A).

ROC analyses: To quantify the firing rate changes, we used an algorithm based on ROC analyses that calculates the ability of an ideal observer to classify whether a given spike rate was recorded in one of two conditions (e.g., during go-cue or no-go-cue odor presentation) (*33*). We defined an auROC equal to 2 (ROCarea – 0.5), with the measure ranging from –1 to 1, where –1 signifies the strongest possible value for one alternative and 1 signifies the strongest possible value for the other.

The statistical significance of these ROC analyses was determined with a permutation test. For this test, we recalculated the ROC curves after randomly reassigning all firing rates to either of the two groups arbitrarily. This procedure was repeated a large number of times (500 repeats for analyses of dynamics (Figs. 2D-E, 3A, 5B-C, S2, S4D-E, S5D-E, and S6D-E), 1000 repeats for all other analyses) to obtain a distribution of values. Subsequently, we calculated the fraction of random values exceeding the actual value. For all analyses, we tested for significance at α = 0.01. Only neurons with a minimum number of three trials for each analyzed condition were included in the analyses.

For analyses of dynamics (width: 100 msec, step: 20 msec), we calculated three measures from the auROC values of correct trials (Figs. 2D, 5B-C, S4D, S5D, and S6D):

1. Time of center of mass: This refers to the time corresponding to the center of mass of the significant points of the auROC values (p < 0.01, permutation test). Only neurons with the significant points for each analyzed condition were included in this analysis.
2. Duration: This refers to the duration in which the auROC values were significant (p < 0.01, permutation test) for five or more consecutive bins, containing the time of center of mass. Only neurons with consecutive bins for each analyzed condition were included in this analysis.
3. Onset time: The onset time refers to the time at which the duration was first evident.

The classification of nLOT neurons: Based on the ROC analyses during the odor-sampling epoch, we classified the nLOT neurons into five types (Figs. 1C and S1). First, we calculated the auROC values of the go-cue versus baseline (1,000 to 0 msec before the end of the inter trial interval) and the no-go-cue versus baseline during the odor-sampling epoch in the correct trials. Based on these values, we defined the cue odor selective population that exhibited significant responses for at least one of the cue odor presentations and cue odor non-selective population (type V neurons). Second, in the cue odor selective population, we also calculated the auROC values of the go-cue versus the no-go-cue during the odor-sampling epoch in the correct trials. Based on these values, we defined go-cue preferred neurons (significant go-cue > no-go-cue, type I neurons) and no-go-cue preferred neurons (significant go-cue < no-go-cue, type II neurons). Finally, in the remaining population, we also calculated the auROC values of cue odors (go-cue + no-go-cue) versus baseline during the odor-sampling epoch in the correct trials. Based on these values, we defined cue excitatory neurons (cue odors > baseline, type III neurons) and cue inhibitory neurons (cue odors < baseline, type IV neurons). For all analyses above, we tested for significance at α = 0.01 (permutation test).

Population vector construction and analyses: We constructed the 2 conditions (71 time bins) × 365 neurons matrix (*34–36*) during the odor-sampling epoch, in which columns contained the auROC values of the correct trials corresponding to the trial-averaged firing rate changes from the baseline (Fig. S7A). By performing principal component analysis (PCA) on the dataset, we reduced the dimensionality of the nLOT population from 365 neurons to three principal components (PCs) and obtained the odor-sampling epoch subspaces. Notably, we used the three subspaces because 82.8% of the total variance was explained (Fig. S7B). To visualize the nLOT population responses, we projected the dataset onto the three-dimensional subspaces (Fig. 3B). This allowed us to obtain a point reflecting the response of the entire population for each of the two conditions at a given instant. The distance between the cue responses was computed as the Euclidean distance between pairs of activity vectors of all subspaces at a given instant (Fig. 3C) (*14, 37*). The velocity of population responses was determined as the distance between successive 20 msec bins (Fig. 3D) (*37*). These values were compared with the values during the baseline epoch (200 to 0 msec before the end of the inter trial interval).

To examine the contribution of individual neurons to the cue encoding, we evaluated the absolute values of PC coefficients as the neural weights (Figs. 3E, S7A, and S7C). We also evaluated contributions along the time course by calculating the absolute values of PC coefficients in the sliding bins (width: 100 msec, step: 20 msec) during odor-sampling (Figs. 3F and S7D). SVM decoding analyses: We used a support vector machine (SVM) algorithm with a linear kernel as a classifier (*14, 15*) and a Matlab function (fitcsvm) for analyses. All analyses were conducted on trial data pooled across animals. A matrix containing concatenated firing rates for each trial and each neuron provided input to the classifier. The matrix dimensions were the number of cells by the number of trials. To avoid over-fitting, k-fold cross-validation (k = 10) was used to calculate the decoding accuracy of trial type discriminations. To compute the decoding accuracy, 40 trials for each trial type (from start of the session) were chosen as the dataset. Next, the dataset was partitioned into ten equal parts—one part was used for testing and the remaining parts were used for training the classifier. This process was repeated ten times to test each individual part; the mean value of the accuracy was used for decoding accuracy. To compute the decoding accuracy of a 100 msec bin window (step: 20 msec), the classifier was trained and tested with a 100 msec bin window (step: 20 msec).

To investigate the relationship between the decoding accuracies and the number of neurons that used them, we calculated the decoding accuracy based on 1-364 randomly sampled neurons (500 repeats) without replacement (Figs. 4 and S8A). Furthermore, we examined the relationship between the decoding accuracy and the contribution of each nLOT neuron type by calculating the decoding accuracy based on 20 randomly sampled neurons (500 repeats) from each neuron group (Fig. 4C). These results were independent of the number of neurons (Fig. S8B).

Statistical analyses: The data were analyzed using MATLAB 2019a. The statistical methods in each analysis have been described in the Results section or figure legends. The Tukey-Kramer method was applied for the tests of significance with multiple comparisons. Although the sample sizes in this study were not pre-determined by sample size calculations, they were based on previous research in the olfactory cortex fields (*15, 38*). Randomization and blinding were not employed. Biological replicates for the histological studies are described in the figure legends.

### Histology

After recording, the mice were deeply anesthetized by an intraperitoneal injection of sodium pentobarbital. Electric lesions were made using 10–20 μA direct current stimulation for 5 sec of one of the four tetrode leads. The mice were perfused transcardially with phosphate-buffered saline (PBS) and subsequently with 4% paraformaldehyde (PFA). The brains were removed from the skull and post-fixed in PFA. Next, the brains were cut into 50-μm thick coronal sections and stained with cresyl violet. The electrode track positions were determined in reference to the atlas developed by Paxinos and Watson (*39*).

## Acknowledgments

We thank Nozomi Fukui for assistance data collection and the lab members for valuable discussions. And we thank Hideki Tanisumi for providing illustrations in the figures. H.M. was supported by the Takeda Science Foundation, and JSPS KAKENHI Grant Numbers 25135708, 16K14557. Y.S. was supported by JSPS KAKENHI Grant Numbers 20H00109, 20H05020.

## Author contributions

Y.T., K.S. and H.M. designed the experiments, Y.T., K.S. and H.M. performed experiments. Y.T., K.S., J.H. and H.M. performed data analysis. Y.T., K.S. and H.M. wrote the paper. Y.S. supported and advised the project.

## Supplementary Materials

**Fig. S1.**
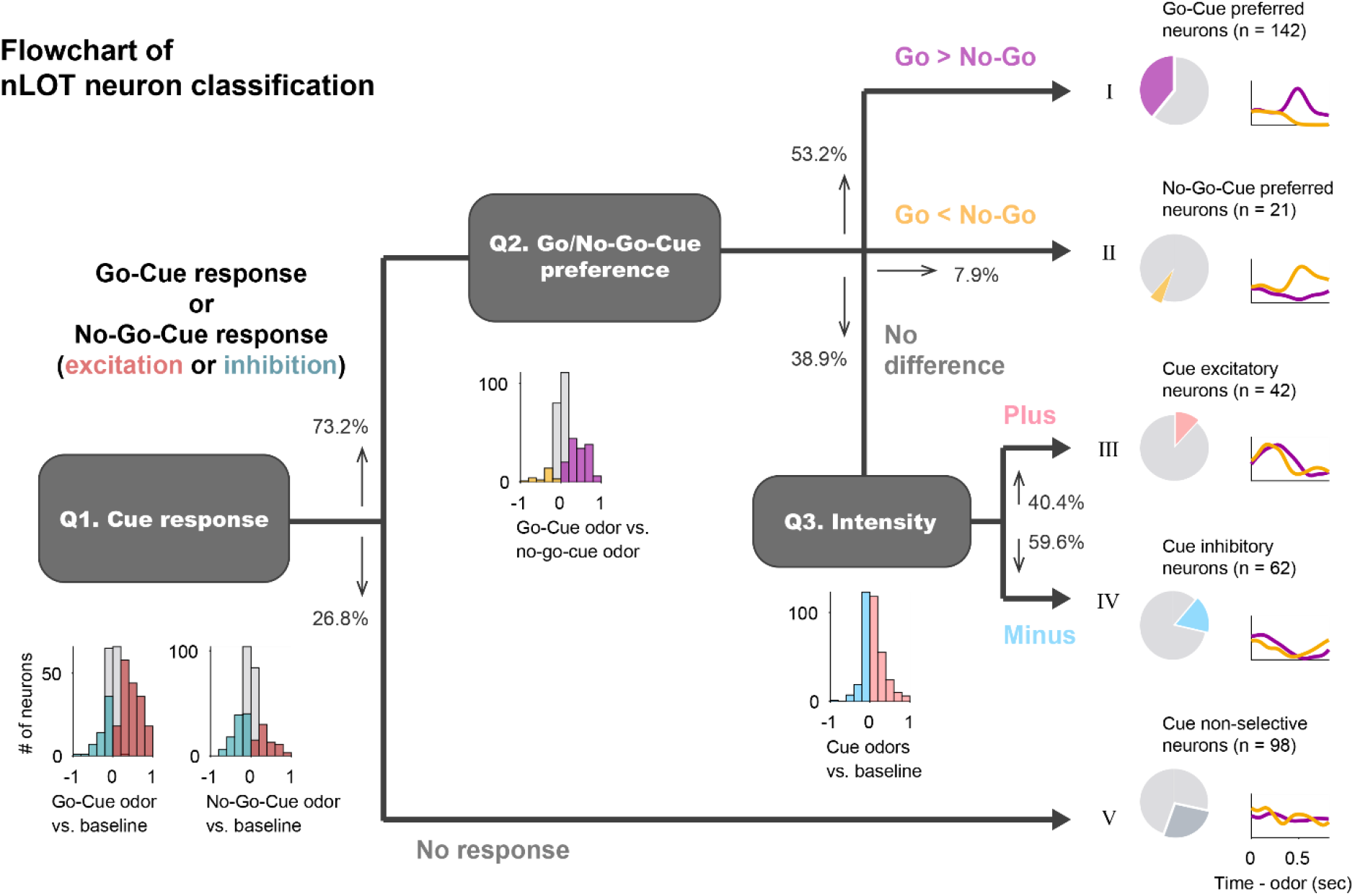
Flowchart of nucleus of the nLOT neuron classification. We classified the nLOT neurons into five types based on the response profiles of the odor-sampling epoch. First (Q1), we calculated the area under the receiver operating characteristic curve (auROC) values of go-cue versus baseline and no-go-cue versus baseline during the odor-sampling epoch in the correct trials (red histogram, significant excitation; blue histogram, significant inhibition). Based on these values, we defined the cue odor selective population (73.2%) that exhibited significant responses for at least one of the cue odor presentations and cue odor non-selective population (26.8%, type V neurons). Second (Q2), in the cue odor selective population, we also calculated the auROC values of go-cue versus no-go-cue during the odor-sampling epoch in the correct trials (purple histogram, significant go-cue > no-go-cue; orange histogram, significant go-cue < no-go-cue). Based on these values, we defined go-cue preferred neurons (53.2%, type I neurons) and no-go-cue preferred neurons (7.9%, type II neurons). Finally (Q3), in the remaining population (38.9%), we calculated the auROC values of cue odors (go-cue + no-go-cue) versus baseline during the odor-sampling epoch in the correct trials (pink histogram, excitation; light blue histogram, inhibition). Based on these values, we defined cue excitatory neurons (40.4%, type III neurons) and cue inhibitory neurons (59.6%, type IV neurons). For all analyses above, we tested for significance at α = 0.01 (permutation test).

**Fig. S2.**
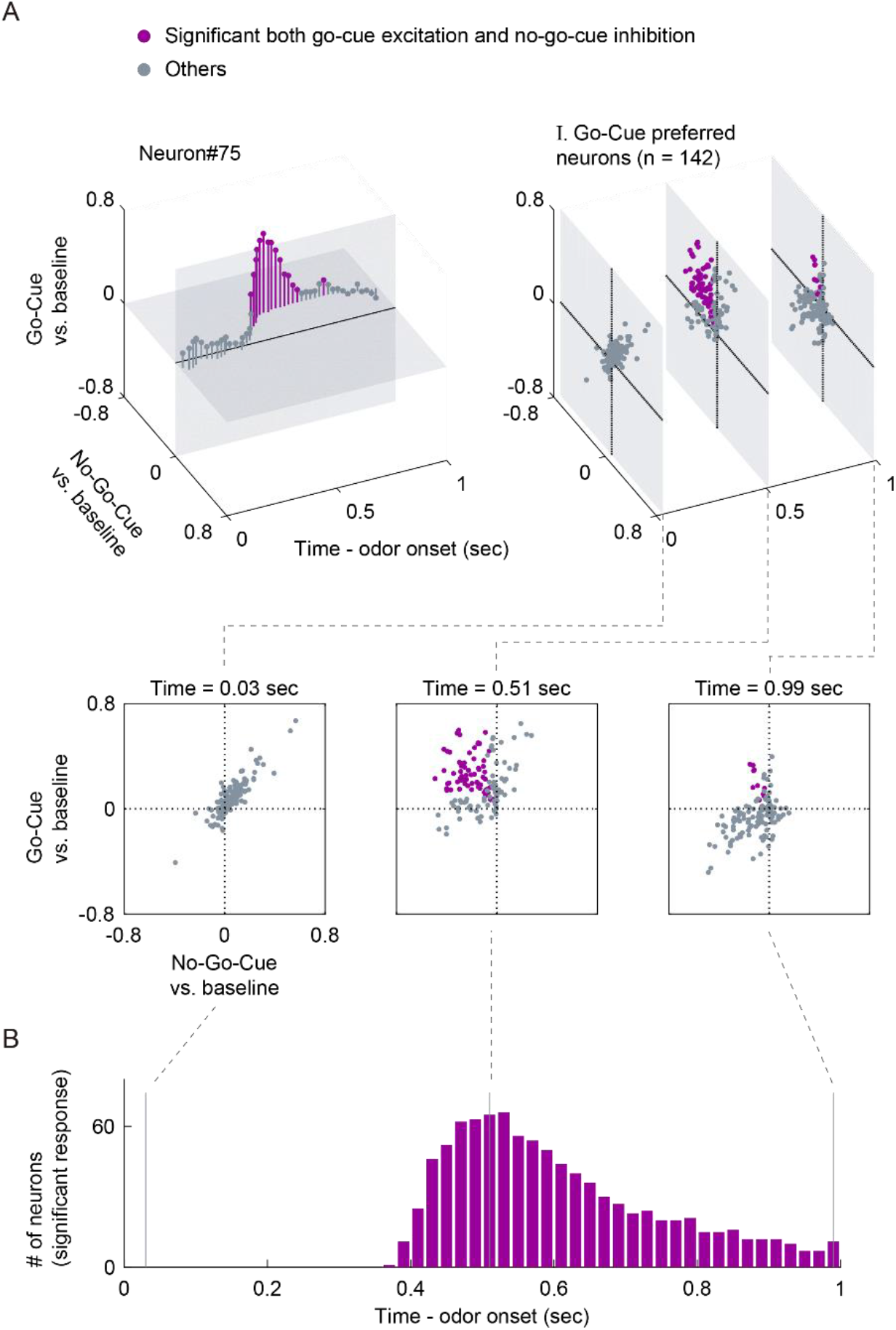
Evaluation of go-cue excitation and no-go-cue inhibition responses. (A) Time course of excitation to go-cue odor and inhibition to no-go-cue odor. Purple dots, significant both go-cue excitation and no-go-cue inhibition (p < 0.01, permutation test); gray dots, other responses. (B) The number of neurons that exhibited significant responses calculated from the area under the auROC values (p < 0.01, permutation test).

**Fig. S3.**
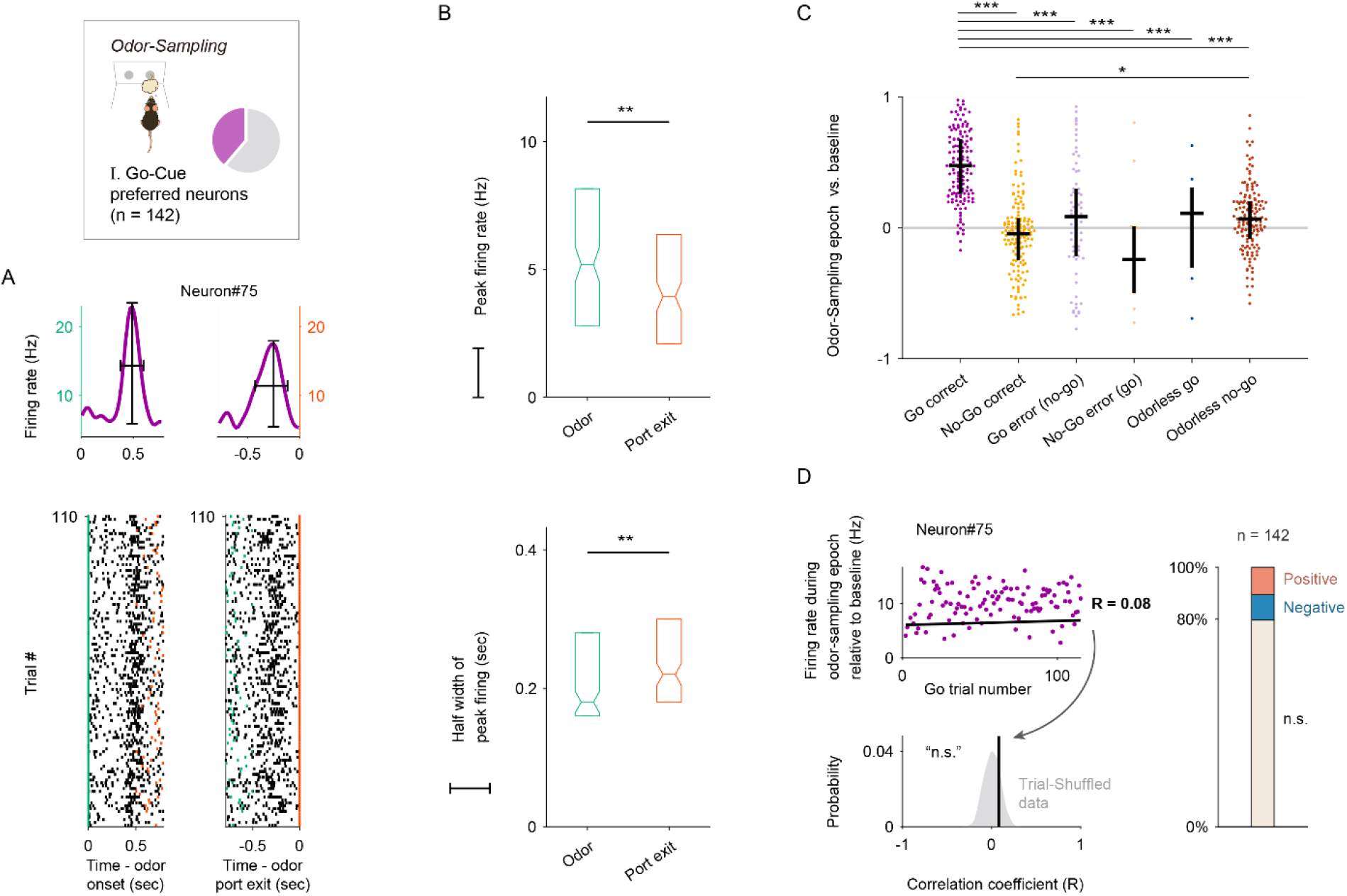
Go-Cue excitation response was triggered by odor onset rather than the initiation of an odor-guided behavior. (A) The activity of an example go-cue preferred neuron aligned to onset of odor valve opening (left, green ticks) or odor port exit (right, red ticks). Raster plots represent the neural activity with each row corresponding to a single trial from the start of the session (bottom) to the 110th trial (top), and each black tick mark to a spike. The peak firing rate (black vertical line) and temporal half width of the peak firing (black horizontal line) were defined from the spike histogram. (B) Comparison of the peak firing rates (top) and half widths of the peak firings (bottom) between the two alignment conditions (odor valve opening versus odor port exit). The peak firing rates were higher when triggered by the odor valve opening (p < 10^-15^, Wilcoxon signed-rank test). Half widths of the peak firings were longer when triggered by the odor port exit (p < 0.001, Wilcoxon signed-rank test). (C) Go-cue excitation and no-go-cue inhibition responses during correct trials, error trials, and catch (odorless) trials. The auROC values were calculated during the odor-sampling epochs and only neurons with a minimum number of three trials for each analyzed condition were included in this analysis. Black horizontal lines and black vertical lines indicate medians and interquartile ranges. The statistical significance among six groups (*p < 0.05, ***p < 0.001) was assessed by one-way analysis of variance (ANOVA) with Tukey’s post hoc test. (D) The development of cue responses in go-cue preferred neurons during learning. For each go-cue preferred neuron, we calculated the correlation between the firing rate during the go-cue odor-sampling epoch relative to the baseline (a mean firing rate during inter trial interval was subtracted for each neuron) and the order of go trial from the start of the session. The correlation coefficient was compared with control values calculated by the 1000 trial-shuffled data (gray shaded area) and then the statistical significance was determined (< 0.5^th^ percentiles of the control values, negative correlation; > 99.5^th^ percentiles of the control values, positive correlation). Across go-cue preferred neurons, the majority of the go-cue responses were not correlated with trial progression (79.5%, not significant; 9.9%, negative; 10.6%, positive).

**Fig. S4.**
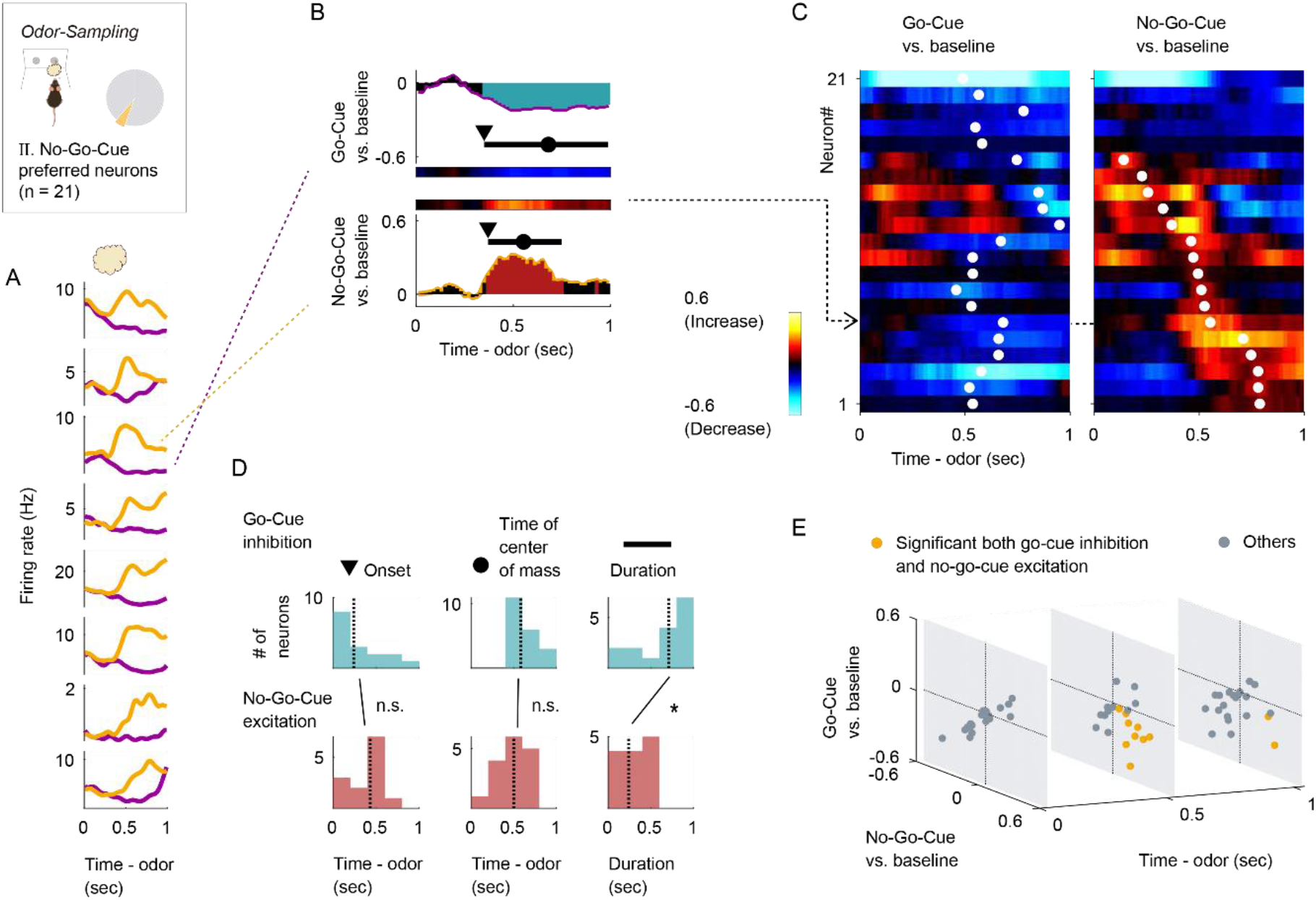
No-Go-Cue preferred neurons showed inhibition to go-cue odor and excitation to no-go-cue odor. (A) Example firing patterns of no-go-cue preferred neurons during the odor-sampling epoch. Spike histograms were calculated using a 20 msec bin width and smoothed by convolving the spike trains with a 60 msec wide Gaussian filter (purple line, go-cue odor; orange line, no-go-cue odor). (B) An example of the area under the auROC values for a no-go-cue preferred neuron. The auROC values (aligned by odor valve opening) were calculated by go-cue odor presentation versus the baseline (top) and no-go-cue odor presentation versus the baseline (bottom) in the sliding bins (width, 100 msec; step, 20 msec). The red bars show significant excitation and blue bars show significant inhibition (p < 0.01, permutation test). Based on the significant time points, the onset times (black triangle), times of center of mass (black circle), and duration (black horizontal line) were calculated. (C) The auROC values for no-go-cue preferred neurons (n = 21, type II neurons). Each row corresponds to one neuron, with neurons in the left and right graphs in the same order. The neurons are sorted by the times of center of mass (white dots) of the auROC values calculated by no-go-cue odor presentation versus the baseline. The color scale is as in (B). An arrow indicates the same neuron as in (B). (D) Distributions of onset time, times of center of mass, and duration for significant inhibitions (top, blue) and significant excitations (bottom, red). Vertical dashed lines indicate the median values. Statistical significance between excitations and inhibitions (*p < 0.05) was assessed by the Wilcoxon rank-sum test. (E) Time course of inhibition to go-cue odor and excitation to no-go-cue odor. Orange dots, significant both no-go-cue inhibition and go-cue excitation (p < 0.01, permutation test); gray dots, other responses.

**Fig. S5.**
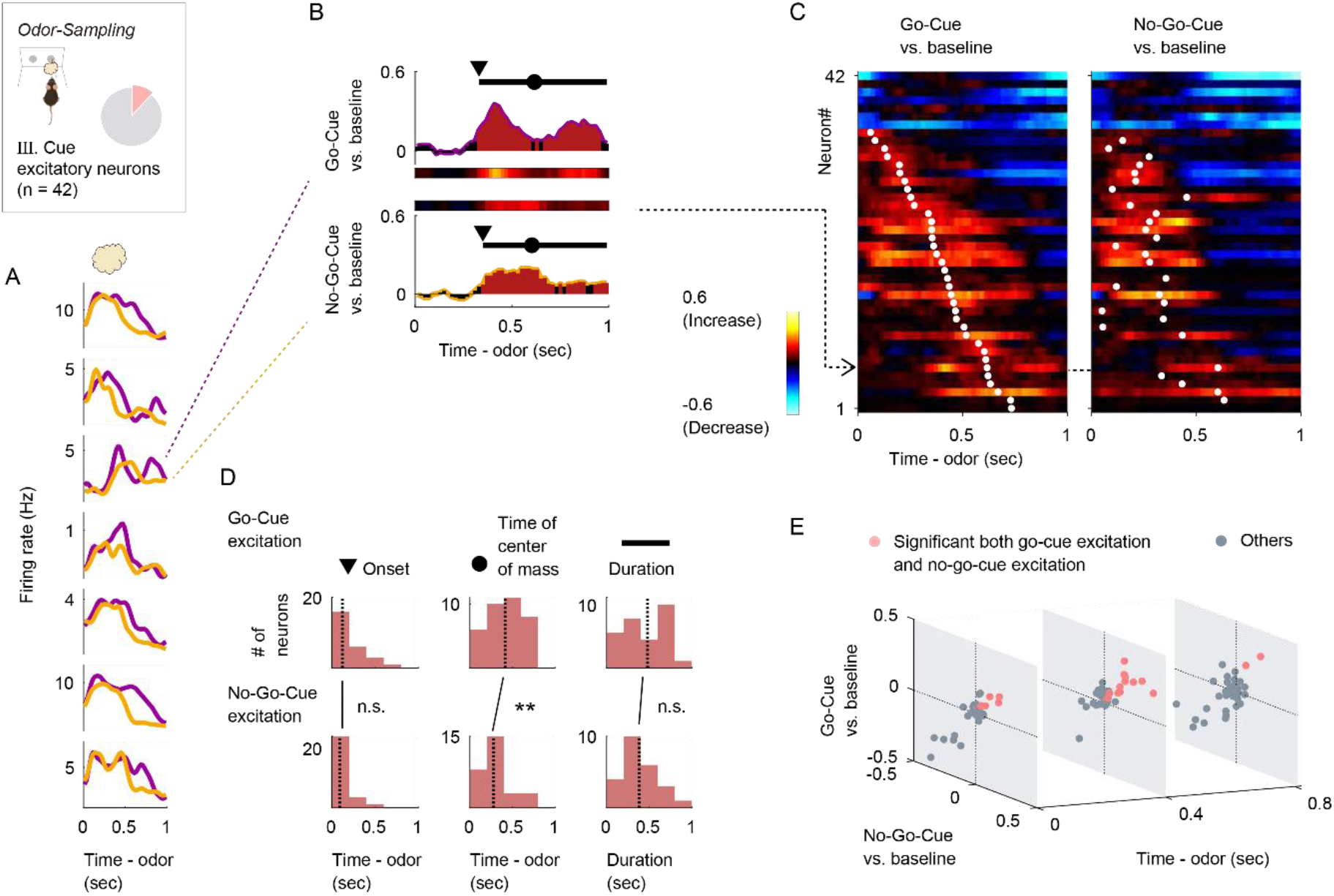
Cue excitatory neuron activity patterns. (A) Example firing patterns of cue excitatory neurons during the odor-sampling epoch. Spike histograms are calculated using a 20 msec bin width and smoothed by convolving spike trains with a 60 msec wide Gaussian filter (purple line, go-cue odor; orange line, no-go-cue odor). (B) An example of the auROC values for a cue excitatory neuron. The auROC values (aligned by odor valve opening) were calculated by go-cue odor presentation versus the baseline (top) and no-go-cue odor presentation versus baseline (bottom) in the sliding bins (width, 100 msec; step, 20 msec). Red bars show significant excitation (p < 0.01, permutation test). Based on the significant time points, onset times (black triangle), times of center of mass (black circle), and duration (black horizontal line) were calculated. (C) The auROC values for cue excitatory neurons (n = 42, type III neurons). Each row corresponds to one neuron, with neurons in the left and right graphs in the same order. Neurons are sorted by times of center of mass (white dots) of the auROC values calculated by go-cue odor presentation versus the baseline. The color scale is as in (B). An arrow indicates the same neuron as in (B). (D) Distributions of onset times, times of center of mass, and duration of for significant excitations. Vertical dashed lines indicate median values. Statistical significance between cue odors (**p < 0.01) was assessed by the Wilcoxon rank-sum test. (E) The time course of excitation to go-cue odor and excitation to no-go-cue odor. Pink dots, significant both go-cue excitation and no-go-cue excitation (p < 0.01, permutation test); gray dots, other responses.

**Fig. S6.**
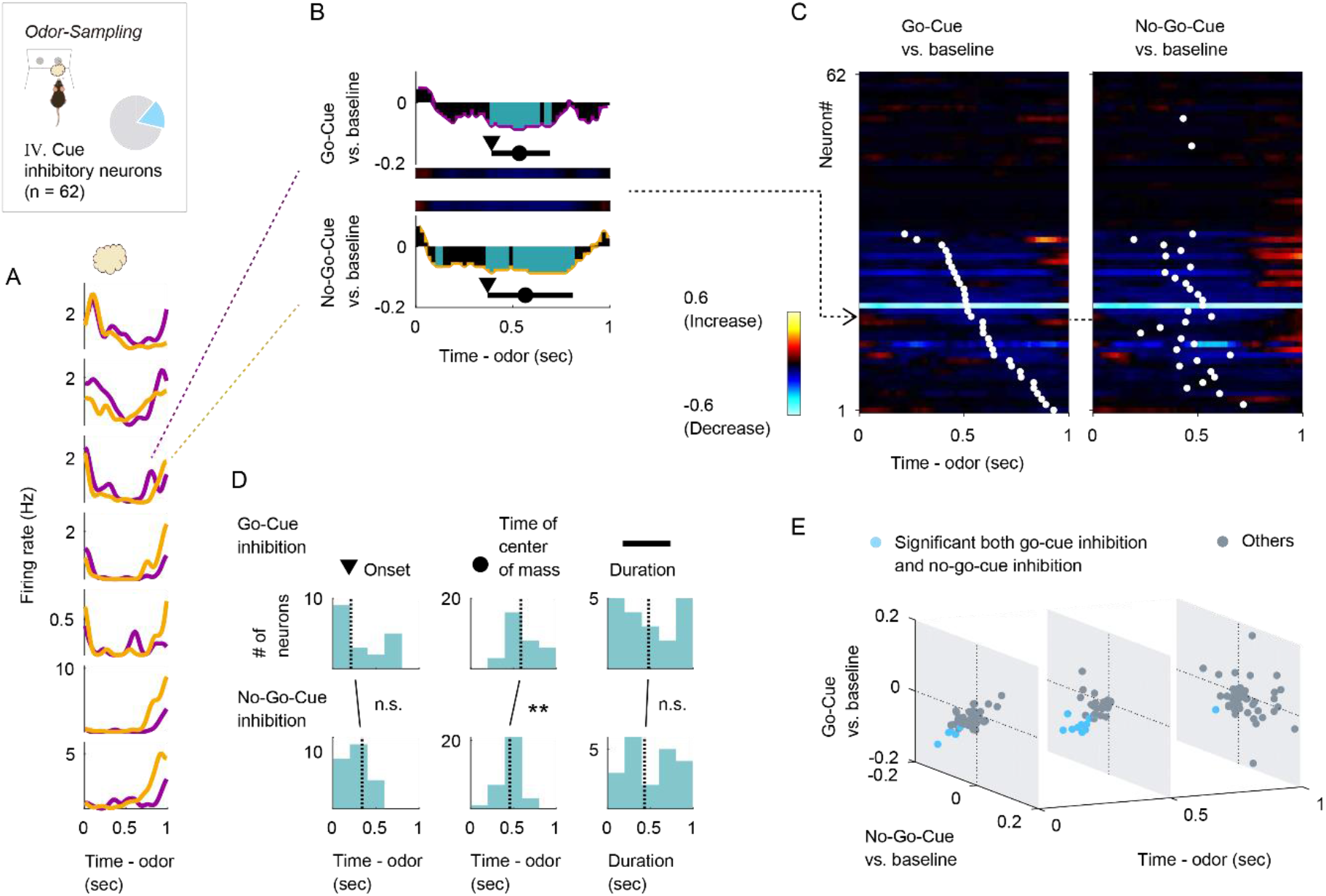
Cue inhibitory neuron activity patterns. (A) Example firing patterns of cue inhibitory neurons during the odor-sampling epoch. Spike histograms are calculated using a 20 msec bin width and smoothed by convolving spike trains with a 60 msec wide Gaussian filter (purple line, go-cue odor; orange line, no-go-cue odor). (B) An example of the auROC values for a cue inhibitory neuron. The auROC values (aligned by odor valve opening) were calculated by go-cue odor presentation versus the baseline (top) and no-go-cue odor presentation versus the baseline (bottom) in the sliding bins (width, 100 msec; step, 20 msec). Blue bars show significant inhibition (p < 0.01, permutation test). Based on the significant time points, onset times (black triangle), times of center of mass (black circle), and duration (black horizontal line) were calculated. (C) The auROC values for cue inhibitory neurons (n = 62, type IV neurons). Each row corresponds to one neuron, with neurons in the left and right graphs in the same order. The neurons are sorted by the times of center of mass (white dots) of the auROC values calculated by go-cue odor presentation versus the baseline. The color scale is as in (B). An arrow indicates the same neuron as in (B). (D) Distributions of onset time, times of center of mass, and duration for significant inhibitions. The vertical dashed lines indicate the median values. The statistical significance between cue odors (**p < 0.01) was assessed by the Wilcoxon rank-sum test. (E) Time course of inhibition to go-cue odor and inhibition to no-go-cue odor. Light blue dots, significant both go-cue inhibition and no-go-cue inhibition (p < 0.01, permutation test); gray dots, other responses.

**Fig. S7.**
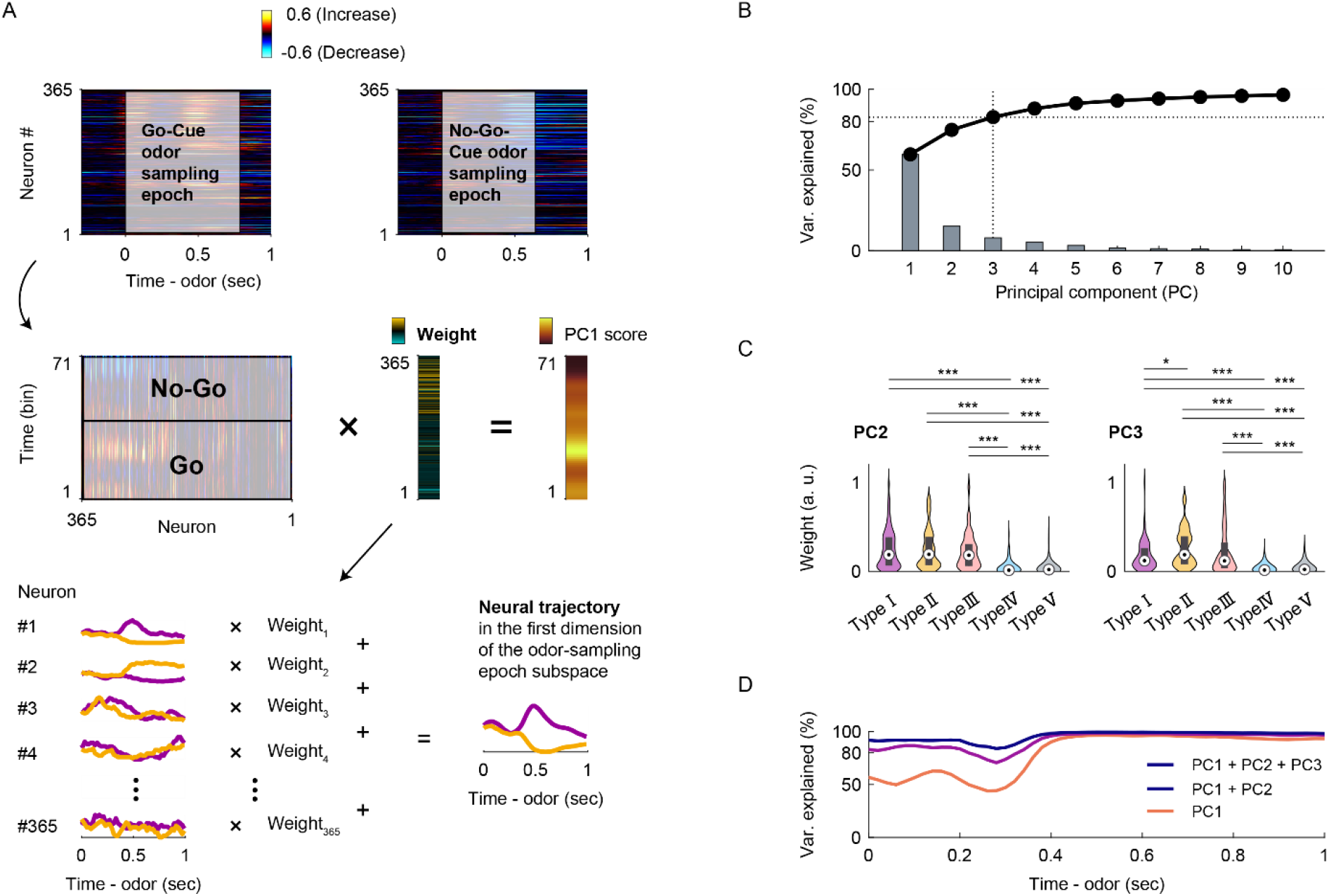
Population vector construction and analyses for the nLOT neuron population response. (A) Population vector construction. We constructed the two conditions (71 time bins) × 365 neurons matrix during the odor-sampling epoch, in which the columns contained the auROC values corresponding to the trial-averaged firing rate changes from the baseline. By performing principal component analysis (PCA) on the dataset, we reduced the dimensionality of the nLOT population from 365 neurons to three principal components (PCs). Subsequently, we obtained the odor-sampling epoch subspaces and neural weights (graphs show the values of the first dimension of the odor-sampling epoch subspaces). (B) Scree plot of the odor-sampling epoch subspaces. It is notable that we used the three subspaces because they explained 82.8% of the total variance. (C) Neural weights in the second (left) and third (right) dimension of the odor-sampling epoch subspaces. Box-plots in violin-plots indicate the medians and interquartile ranges. Purple, type I; orange, type II; pink, type III; light blue, type IV; gray, type V. Statistical significance among five groups (*p < 0.05, ***p < 0.001) was assessed by one-way analysis of variance (ANOVA) with Tukey’s post hoc test. (D) Variances of neural weights data along the time course (Fig. 3F) in the dimensions of each sliding bin (width: 100 msec, step: 20 msec).

**Fig. S8.**
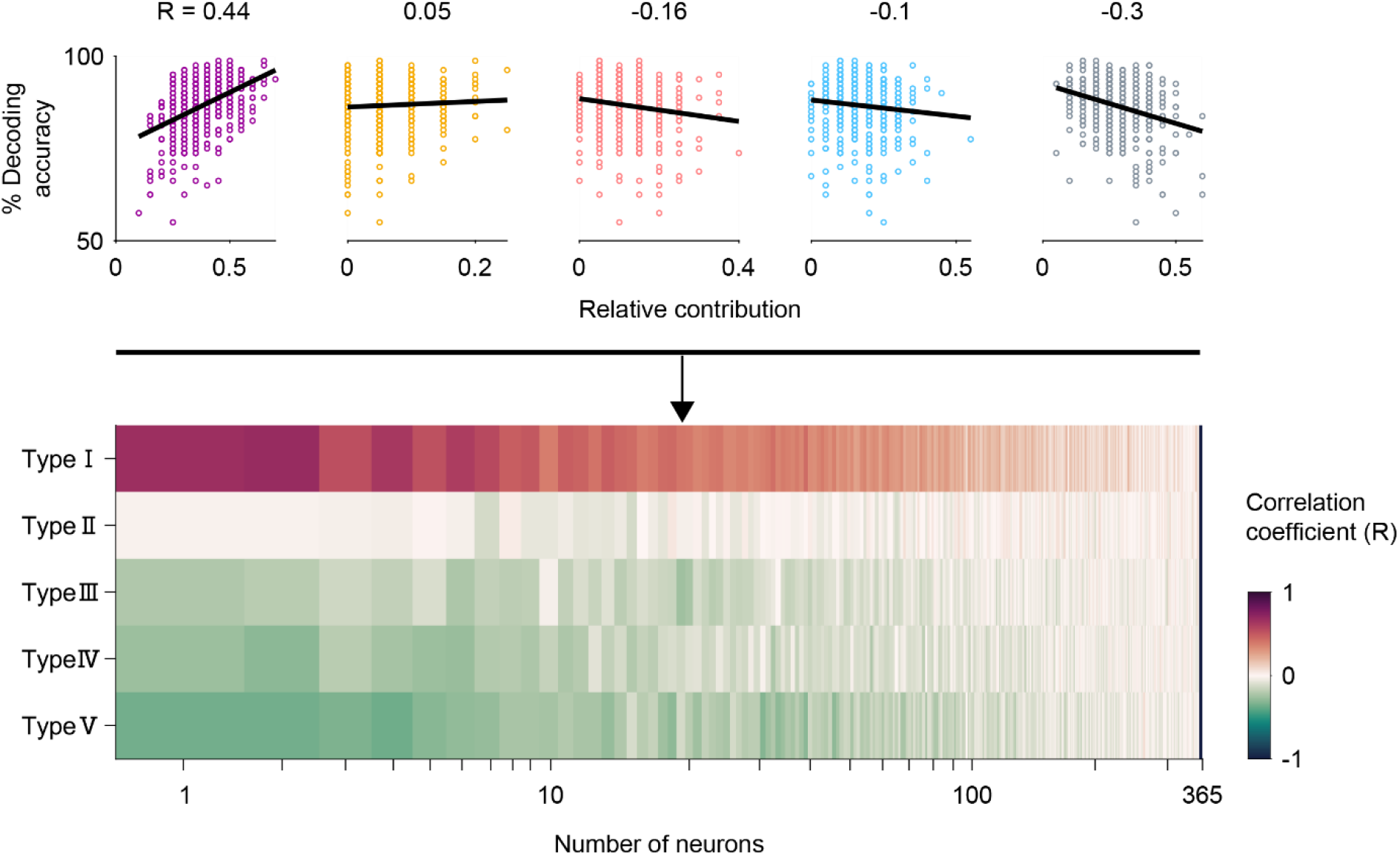
Decoding accuracies based on 1–364 randomly sampled neurons during the 400–500 msec period after the odor onset. Correlations between the decoding accuracies and relative contributions as the proportion of each neuron type number in randomly sampled neurons. The decoding accuracies based on 1–364 randomly sampled neurons without replacement during the 400–500 msec period after the odor onset.

**Fig. S9.**
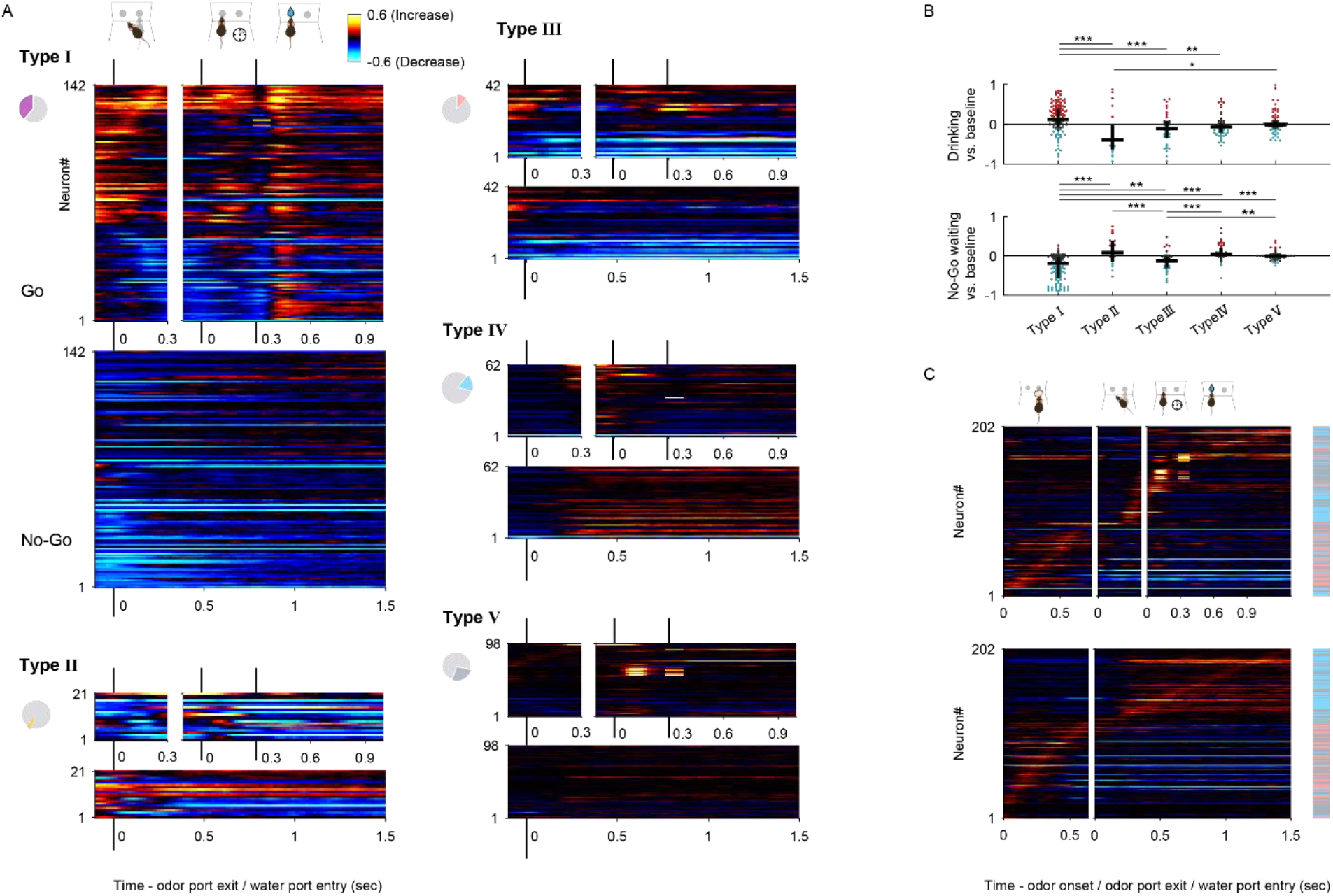
Response profiles following odor-guided behaviors. (A) The auROC values calculated by go or no-go trials versus the baseline in the sliding bins (width, 100 msec; step, 20 msec) following odor-guided behaviors. Each row corresponds to one neuron, with neurons in all the graphs in the same order for each neuron group. Neurons were sorted by the peak time for the auROC values. The color scale is as in Fig. 2C. (B) The auROC values during the drinking epoch (top) and no-go waiting epoch (bottom). Black horizontal lines and black vertical lines indicate the medians and interquartile ranges. Red dots, significant excitation; blue dots, significant inhibition; gray dots, non-significant (p < 0.01, permutation test). Statistical significance among five groups (*p < 0.05, **p < 0.01, ***p < 0.001) was assessed by one-way analysis of variance (ANOVA) with Tukey’s post hoc test. (C) The auROC values calculated by go or no-go trials versus the baseline in the sliding bins (width, 100 msec; step, 20 msec) during odor-guided go/no-go task in the type III, IV, and V neurons. Each row corresponds to one neuron. Neurons are sorted by the peak time for the auROC values. The color scale is as in Fig. 2C. The colored box on the right shows neuron type for each neuron (pink, type III; light blue, type IV; gray, type V). Note that these neurons tended to show an excitatory response to a specific behavioral epoch with inhibitory responses relative to other behavioral epochs.

